# GlycoBind: a general toolkit for protein-glycan binding prediction

**DOI:** 10.64898/2026.11.25.769100

**Authors:** Xirui Huang, Zhixin Tian

**Affiliations:** School of Chemical Science & Engineering, Shanghai Key Laboratory of Chemical Assessment and Sustainability, Tongji University, Shanghai 200092, China

**Keywords:** protein-glycan interaction, glycan array, graph representation, binding prediction, candidate prioritization

## Abstract

Protein-glycan recognition controls immune responses, host-pathogen adhesion, cell signalling and the functional organization of the glycocalyx. Computational models can predict protein binding to monosaccharides or short oligosaccharides, but predicting binding to structurally diverse intact glycans remains challenging. Here we present GlycoBind, a general toolkit that accepts a protein structure in Protein Data Bank (PDB) format and an International Union of Pure and Applied Chemistry (IUPAC)-condensed glycan, and returns a binding score without requiring a protein-glycan complex or a docking pose. GlycoBind adopts the frozen residue-state representation of DeepGlycanSite, a carbohydrate-binding-site predictor, and combines this protein representation with frozen GlycanGT graph representations through a compact trainable classifier. Across five protein-grouped holdouts, GlycoBind achieved a mean area under the receiver operating characteristic curve (AUROC) of 0.767. On six external glycan-array datasets after exact protein and glycan cleaning for both models, GlycoBind showed higher aggregate binding-strength ranking performance and stronger recovery of high-intensity binding pairs than the existing models. By prioritizing high-intensity pairs for experimental follow-up, GlycoBind may support mechanistic studies of protein-glycan recognition and the rational development of glycan-targeted therapeutics.

## Introduction

Protein-glycan recognition is central to biological regulation. Glycans participate in protein folding and stability, cell-cell communication, immune recognition and the organization of the cell-surface glycocalyx ^1^. In immunity, lectins decode glycan structures to distinguish self from non-self and to regulate inflammatory responses ^2-3^. Glycan recognition also determines host-pathogen adhesion and tissue tropism during infection ^4^. These biological roles have made protein-glycan interfaces attractive targets for vaccines, anti-adhesive agents and glycomimetic drugs directed at selectins, Siglecs and other lectins ^5^. High-throughput glycan arrays and associated analysis tools provide a scalable way to survey this interaction space ^6-8^, but the resulting data are sparse, imbalanced and heterogeneous across proteins and array formats. Computational prioritization is therefore needed before candidate pairs advance to lower-throughput biochemical or structural experiments.

Learned protein-carbohydrate scoring has advanced rapidly for monosaccharide and short- oligosaccharide settings. LectinOracle-SweetNet, GlyNet and CSM-carbohydrate have shown strong performance in this regime by combining protein-language representations, glycan-array multitask learning or structure-informed affinity scoring ^9-11^. Machine-learning annotation of a large lectin panel has further improved the interpretation of glycan-array binding profiles ^12^, whereas broad reviews of computational tools and array-analysis workflows now connect measurement, motif analysis and prediction ^7-8,13^. These studies establish that compact glycan inputs can support accurate predictions when informative training measurements are available. Graph-based learning has also introduced transferable representations for branched glycans ^14^, including large-scale analysis of glycan 3D structures ^15^ and pretrained graph-transformer representations for complete glycan structures ^16^. For protein-side localization, DeepGlycanSite predicts carbohydrate-binding residues directly from protein structures ^17^, and structure-based neural models have extended protein-carbohydrate interaction prediction from residue-level contacts to proteome-scale inference ^18-19^.

MCNet further advances protein-glycan modelling toward atom-level interaction prediction ^20^. A practical pair-level toolkit must still combine a protein structure, a complete glycan graph and an explicit binding score in one reproducible workflow. Recent graph-based and structure-based protein-carbohydrate models have extended pair classification, binding-site and binding-interface prediction ^21-26^. Broader reviews of glycoinformatics and protein-carbohydrate modelling provide context for these developments ^27-28^, while public glycan-array resources support data reuse and audit ^29^.

Extending this task from monosaccharides and short oligosaccharides to large branched glycans introduces additional computational costs. Glycans remain conformationally dynamic and often act through multivalent presentation ^30-31^. Glycosidic linkages create large conformational search spaces, and docking must sample both glycan internal conformations and protein-interface poses ^32^.

Specialized methods such as Vina-Carb refine glycosidic angles during carbohydrate docking ^33^; GlycanDock supports glycoligand docking and refinement in Rosetta ^34^; HADDOCK provides an integrative workflow for modelling protein-glycan interactions ^35^; and GlycoTorch Vina extends Vina-based docking to glycosaminoglycans ^36^. Recent water-guided docking has further improved prediction of protein-glycan complexes ^37^. These tools are valuable when a binding pose is the experimental objective. However, reported docking accuracy and confidence generally decrease for larger and more branched glycans as the search space expands ^32-35,37^. Pose ranking also remains sensitive to sampling and scoring, and many established scoring functions are formulated for an already assembled protein-glycan complex ^11,35-37^. Pair-level screening therefore needs a complementary entry point that can score a protein structure and glycan sequence without docking every candidate.

We developed GlycoBind to meet this need. The toolkit adopts the frozen residue-state matrix generated inside DeepGlycanSite as a general protein representation and uses its site probabilities only to weight a residue summary. These protein features are paired with frozen GlycanGT representations of the intact IUPAC-condensed glycan before a compact fusion and classification head is trained. We evaluated the model with five protein-disjoint holdouts, a retrained feature-branch ablation, recovery of high-intensity pairs and six external glycan-array datasets. External comparisons used identical task rows and removed exact protein and exact glycan overlap with both GlycoBind and the released LectinOracle-SweetNet model. The resulting evidence supports GlycoBind as a tool for binding-strength ranking and candidate prioritization of protein-glycan interactions, rather than a substitute for pose prediction, affinity measurement or experimental validation.

## Experimental section

### Source data and binding labels

The original glycan-array dataset was organized and released by the LectinOracle authors^9^. We used their processed matrix as the starting point. Protein sequences were mapped to stable identifiers and AlphaFold2/ColabFold structures ^38^. Array responses were treated as normalized z-score-like measurements. Following the one-sided glycan-array binding rule of Cholleti et al. ^39-40^, pairs with z>1.0364 were labelled positive, corresponding to p<0.15, and pairs with z<0 were labelled negative. Pairs with 0<=z<=1.0364 were treated as a gray zone and excluded from fitting and evaluation.

The final labelled benchmark contained 382,782 protein-glycan pairs, including 30,942 positive and 351,840 negative pairs, and covered 1,141 proteins and a graph library of 927 glycans.

Positive prevalence was only 8.1%, so plain accuracy was dominated by the negative class and was not used to rank models. AUROC and the area under the precision-recall curve (AUPRC) were treated as the primary threshold-free measures, and balanced accuracy and Matthews correlation coefficient (MCC) were used to characterize operating-point and ranking behavior.

### Protein structures and DeepGlycanSite intermediate representations

All development proteins were modelled with AlphaFold2/ColabFold using multiple sequence alignments (MSAs) as input and stored as PDB files ^38^. The selected chain must match the reference sequence with at least 50% identity. Structures shorter than 30 amino acids or with mean C-alpha predicted local distance difference test values below 70 were excluded.

The receptor-only DeepGlycanSite model was run with frozen weights ^17^. Protein residues combined amino-acid identity, local geometry, backbone and side-chain torsions and layer-33 ESM-2 650M embeddings ^38^. Layer 33 is the final representation of the 33-layer ESM-2 650M encoder. The residue graph was processed by an 11-layer equivariant network and a receptor-only site decoder. For a protein of length L, the outputs were an Lx512 residue-state matrix and an L-dimensional site-probability vector.

DeepGlycanSite was developed for carbohydrate-binding-site prediction. Here, its final site classification was not used as the output of GlycoBind. Instead, the Lx512 residue-state matrix was retained as a frozen intermediate protein representation, and the site probabilities were used only to weight the top-24 residue summary.

### Glycan graphs and GlycanGT representations

IUPAC-condensed glycans were converted to monomer-linkage graphs. Each graph contained monomer nodes and linkage relations. The tokenizer and vocabulary were compatible with GlycanGT ^16^. GlycanGT was run with frozen weights and returned a 768-dimensional graph token and an Nx768 matrix of monomer-node tokens. Linkage tokens participated inside GlycanGT self-attention.

### Model architecture and training objective

At inference, a protein structure in PDB format enters the frozen receptor-only DeepGlycanSite model, which produces an Lx512 residue-state matrix and an L-dimensional site-probability vector. In parallel, the IUPAC-condensed glycan is parsed into a monomer-linkage graph and passed through the frozen GlycanGT encoder, which returns one 768-dimensional graph token and N node tokens of 768 dimensions. Only the downstream projection, fusion and classification layers are trainable; the DeepGlycanSite and GlycanGT parameters remain fixed throughout fitting.

The protein branch used the mean of all residue states and a probability-weighted sum of the 24 residues with the highest DeepGlycanSite site probabilities. Each 512-dimensional vector was projected to 256 dimensions with a linear layer, LayerNorm and GELU. The vectors were concatenated and compressed to a 256-dimensional protein representation.

The glycan branch projected the graph token from 768 to 256 dimensions with LayerNorm and GELU. Masked mean and masked maximum pooling of node tokens produced a 1,536-dimensional vector, which was projected to 256 dimensions. The two 256-dimensional glycan views were concatenated and compressed to a 256-dimensional glycan representation. The protein and glycan vectors were concatenated into a 512-dimensional feature and passed through 512→128→32→1 layer. A sigmoid transformed the final logit into a binding probability. Each trainable model contained 1,188,801 parameters; DeepGlycanSite and GlycanGT remained frozen.

The training objective was binary cross-entropy (BCE) with logits. The positive-class weight was the square root of the negative-to-positive ratio. Models were optimized with Adam using a learning rate of 3×10^−5^, weight decay of 1×10^−4^, batch size 1,024, dropout 0.3, a maximum of 50 epochs and early stopping after five epochs without improvement in validation AUROC. The epoch with the highest validation AUROC was retained.

### Protein-grouped splits, calibration and decision rule

To evaluate training and evaluation stability, five random protein-grouped partitions (seeds 42-46) were generated for the labelled benchmark. The benchmark contained 1,141 proteins. Exact duplicate protein sequences were assigned to one group before partitioning. For each partition, 80% of protein groups formed the training pool, 10% were used for validation and 10% as an outer test. The complete training pool was used for gradient updates. Validation proteins were used for early stopping and Platt scaling, while outer-test proteins were excluded from training, early stopping, probability calibration and model selection. No exact protein sequence was shared between training, validation and outer-test partitions.

Validation-set Platt scaling was used to display probability calibration. For each seed, an operational raw-probability threshold was selected on the validation set at an empirical false-positive rate of 15%, choosing the threshold with the highest sensitivity among candidates satisfying that constraint. The five thresholds were 0.3360, 0.2986, 0.2877, 0.3086 and 0.2906 for seeds 42-46. Their arithmetic mean, 0.3043, was used as the unified threshold for external inputs. A new protein-glycan pair was classified as binding when the mean of the five raw member probabilities was at least 0.3043.

### Evaluation metrics

Threshold-free metrics included AUROC, AUPRC, protein-macro AUROC, protein-macro Spearman correlation and normalized discounted cumulative gain at the top 10% of each protein profile (NDCG@10%). Thresholded metrics included BACC and MCC. After all model scores were fixed, BACC and MCC were summarized at empirical false-positive rates of approximately 15% for descriptive operating-point comparison; these post hoc operating-point summaries were not used for model fitting or model selection. Probability calibration was assessed by the Brier score and expected calibration error (ECE).

For internal NDCG@10%, each protein profile was evaluated separately with k=max(1, ceiling(0.10n)). Binary binding labels were used as relevance. Discounted cumulative gain used gains of 2^relevance - 1 and logarithmic position discounts. The ideal ranking was obtained by sorting the same tasks by experimental relevance. Internal NDCG therefore measures recovery of binary binding labels, whereas the external NDCG uses the continuous experimental response percentile as graded relevance.

### Benchmark composition

The protein-family and glycan structural-class composition of the labelled benchmark are reported **(Figure 1a,b)**. The workflow from PDB and IUPAC input to a binding score is illustrated **(Figure 1c)**. The Model architecture and training objective section describes the full processing path, frozen components and dimensional changes.

**Figure 1.**
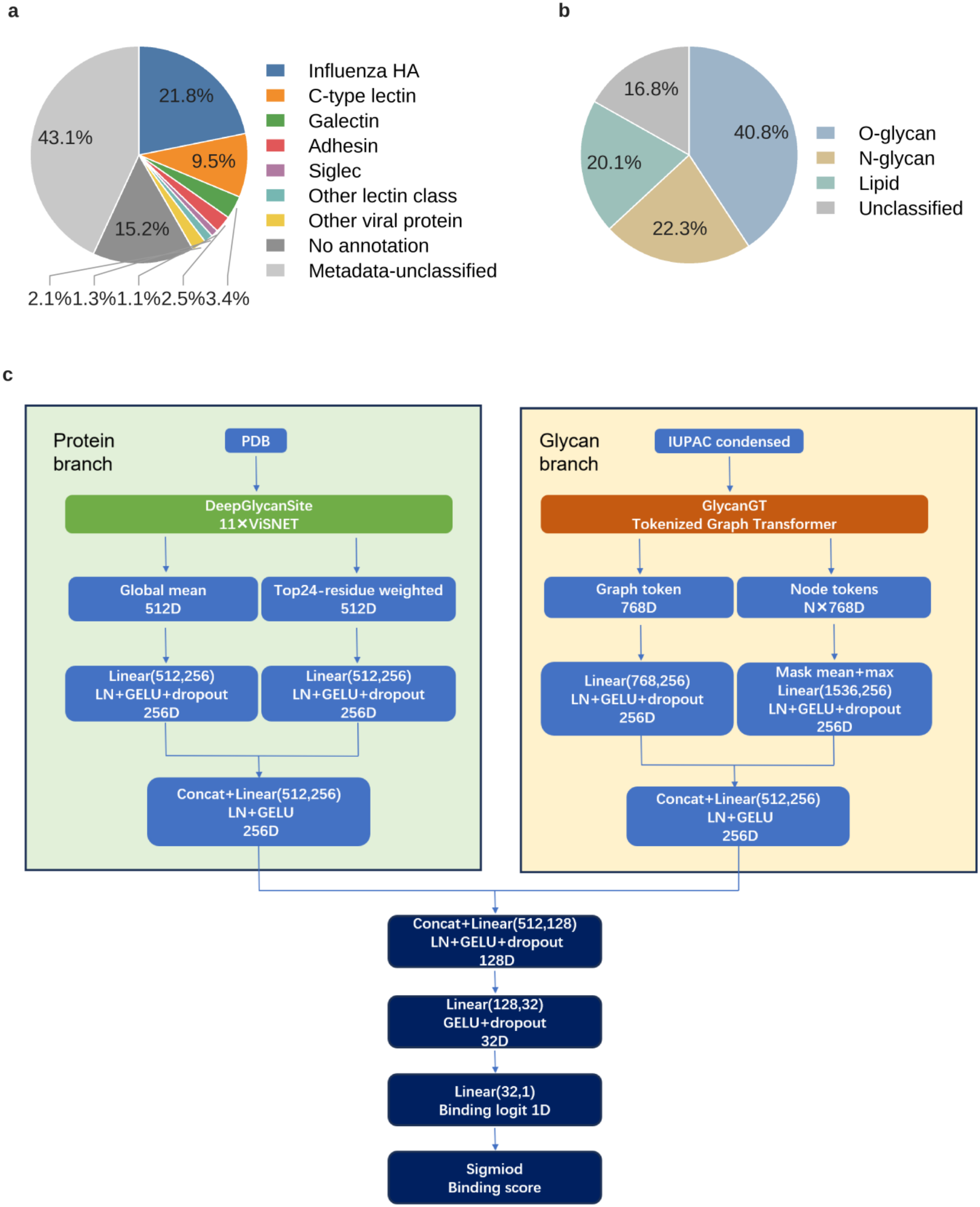
Benchmark composition and model architecture. (a) protein-family composition of the labelled benchmark; (b) glycan structural-class composition; (c) workflow from a protein structure in PDB format and an IUPAC-condensed glycan to a binding score; LN, layer normalization; GELU, Gaussian Error Linear Unit, a smooth neural-network activation function.

### Five-seed discrimination and candidate enrichment

Across seeds 42-46, GlycoBind achieved a mean AUROC of 0.767±0.012 standard deviation (s.d.) and an AUPRC of 0.314±0.023. At post hoc FPR -oriented operating points of approximately 15%, BACC was 0.693±0.009 and MCC was 0.270±0.005. Protein -macro AUROC was 0.751±0.006, protein-macro Spearman correlation was 0.223±0.013 and NDCG@10% was 0.451±0.018. The five - seed receiver operating characteristic curves and the complete metric set are reported **(Figure 2a; Table 1)**.

**Table 1.** Five-seed protein-held-out performance. Values are mean±sample s.d. across seeds 42 -46. BACC and MCC are descriptive operating-point summaries at empirical FPR levels of approximately 15% and were not used for model selection. AUROC, area under the receiver operating characteristic curve; AUPRC, area under the precision-recall curve; BACC, balanced accuracy; MCC, Matthews correlation coefficient; NDCG@10%, normalized discounted cumulative gain at the top 10% of each protein profile; ECE, expected calibration error.

| **Metric** | **Mean±s.d.** |
|---|---|
| AUROC | 0.767±0.012 |
| AUPRC | 0.314±0.023 |
| BACC | 0.693±0.009 |
| MCC | 0.270±0.005 |
| Protein-macro AUROC | 0.751±0.006 |
| Protein-macro Spearman correlation | 0.223±0.013 |
| NDCG@10% | 0.451±0.018 |
| Brier score | 0.0625±0.0054 |
| ECE | 0.0085±0.0011 |

**Figure 2.**
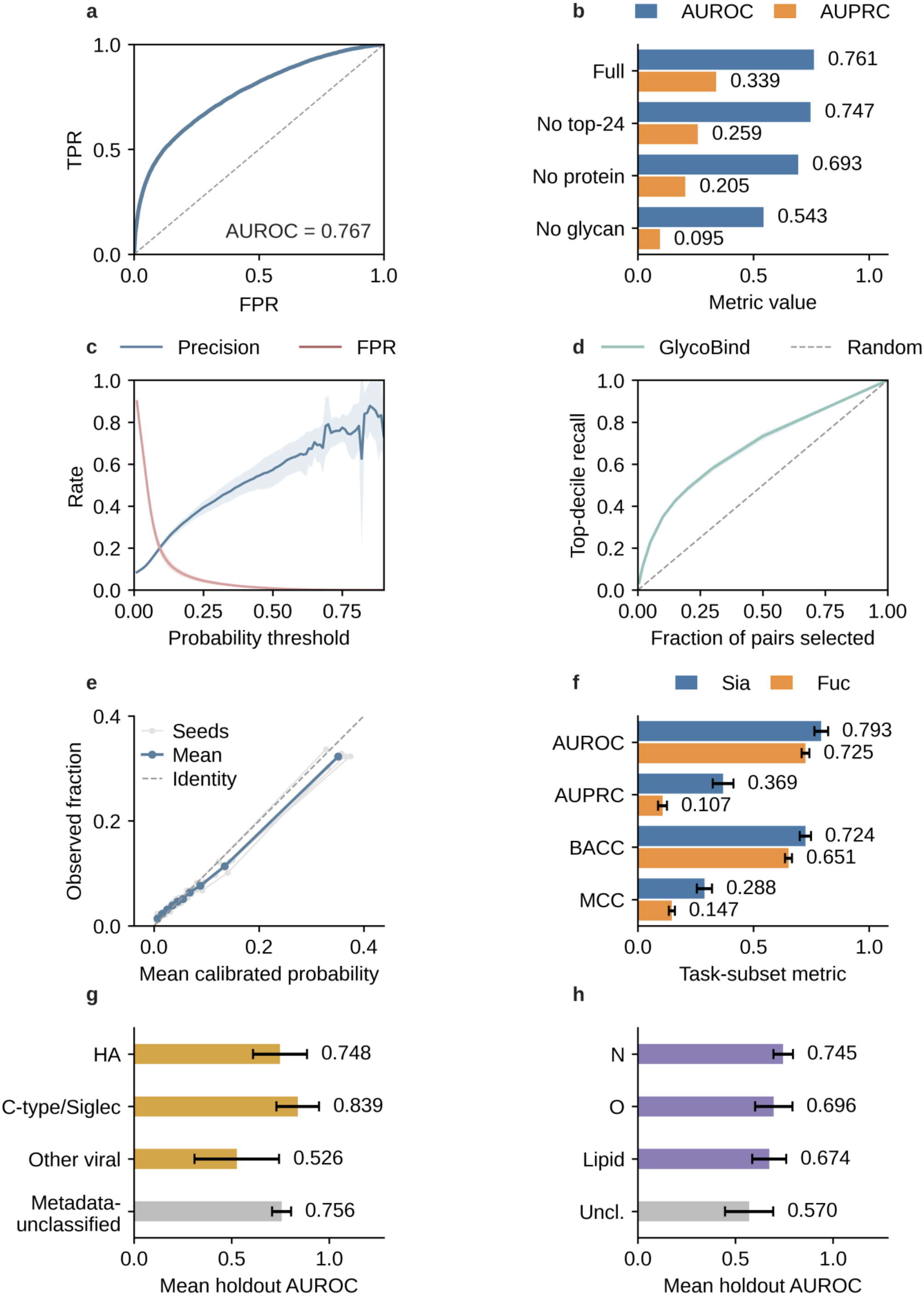
Model evaluation and stratified performance. (a) receiver operating characteristic curves across seeds 42-46; (b) retrained feature-branch ablation comparing AUROC and AUPRC; (c) precision and false-positive rate across probability thresholds; (d) recovery of experimentally high-intensity protein-glycan pairs relative to random selection; (e) probability calibration across bins; (f) Sia and Fuc terminal-subset performance across AUROC, AUPRC, balanced accuracy and MCC; (g) holdout AUROC by protein family; (h) holdout AUROC by glycan class. Error bars and shaded ranges indicate s.d. across seeds unless stated otherwise. FPR, false-positive rate; TPR, true-positive rate; AUPRC, area under the precision-recall curve; MCC, Matthews correlation coefficient.

The mean false-positive rate at the descriptive operating points was 14.98%, with mean sensitivity 53.68%, specificity 85.02% and precision 23.22%. The predicted positive fraction was 17.99%. Platt-calibrated probabilities had a Brier score of 0.0625±0.0054 and an ECE of 0.0085±0.0011. The precision and false-positive rate at each probability threshold and calibration across probability bins are reported **(Figure 2c,e)**.

To quantify recovery of high-intensity responses, the experimental top decile was defined within each protein profile. These top-decile indicators were then pooled across proteins, and recovery was computed from the global ranking of model scores. Selecting the top 10% of pooled pairs recovered 34.9±0.8% of all experimental top-decile pairs, corresponding to 3.49-fold enrichment over random selection. Selecting the top 5% recovered 22.8±0.8% of the experimental top -decile pairs. Recovery over the full selection range is reported **(Figure 2d)**. This is a pooled ranking analysis and should not be interpreted as a per-protein calibration measure.

### Retrained ablation identifies the dominant information sources

Retrained feature-branch ablations were performed on seed 42. The full model achieved AUROC 0.761, AUPRC 0.339, BACC 0.692 and MCC 0.252. Removing the top-24 residue summary changed AUROC to 0.747 and AUPRC to 0.259. The effect of each ablation on AUROC and AUPRC is reported **(Figure 2b)**. Removing the protein branch reduced AUROC to 0.693 and AUPRC to 0.205, and removing the glycan branch reduced AUROC to 0.543 and AUPRC to 0.095.

Within this single split, the glycan graph produced the largest reduction in discrimination, while the protein branch contributed complementary information. Removing the top-24 summary reduced AUROC by 0.014 and AUPRC by 0.079, indicating that the local residue view contributes more clearly under the full training-pool protocol.

### Sialic acid and fucose subset performance

The same model also ranked terminal-subset evaluation rows containing sialic acid (Sia) or fucose (Fuc) without task-specific fine-tuning. For each parent-child glycan pair, the score of the intact glycan was compared with that of the corresponding glycan lacking the terminal Sia or Fuc; the score difference was used to identify proteins whose prediction depended on that terminal sugar or at least interacted with it. For Sia, AUROC was 0.793±0.029, AUPRC was 0.369±0.045, BACC was 0.724±0.024 and MCC was 0.288±0.033. For Fuc, AUROC was 0.725±0.017, AUPRC was 0.107±0.019, BACC was 0.651±0.015 and MCC was 0.147±0.013. The thresholded subset values use the same full-benchmark descriptive FPR-oriented operating point; AUROC and AUPRC are the primary subset comparisons. The Sia and Fuc subset results and the holdout AUROC by protein family and glycan class are reported **(Figure 2f-h)**.

Performance was not uniform across protein families or glycan classes. For protein-family groups represented in at least three holdout seeds, mean AUROC was 0.839±0.109 for C -type lectin/Siglec proteins, 0.756±0.048 for metadata-unclassified proteins, 0.748±0.138 for influenza HA and 0.526±0.217 for other viral proteins. For glycan classes, mean AUROC was 0.745±0.050 for N - glycans, 0.696±0.096 for O-glycans, 0.674±0.087 for lipid or free glycans and 0.570±0.124 for unclassified structures. These group sizes are unequal and the comparison is descriptive, but it identifies C-type lectin/Siglec proteins and N- and O-glycans as the strongest regions of the current applicability domain, whereas other viral proteins and unclassified glycan structures approached chance performance. Rare protein classes are retained in the benchmark composition but omitted from the stratified analysis because they do not meet the three-seed reporting threshold **(Figure 1a; Figure 2g)**. The stratified AUROC results for the remaining groups are reported **(Figure 2g,h)**.

## External validation

### External array datasets and exact cleaning

Six external datasets were used: Rotavirus B VP8*B ^41^, Sasmal AB5 ^42^, EclA ^43^, Influenza H5 ^44^, bisected N-glycan ^45^ and a LectinOracle-SweetNet main-figure subset ^9^. Continuous array responses were analysed by within-profile ranking. The binary Sasmal panel was additionally evaluated with AUROC and AUPRC.

The six external datasets correspond to Tests I-VI in the order listed above **(Figure 3)**. After exact cleaning, Test I retained the single Rotavirus B VP8*B protein, with 3.6/4.7% maximum sequence identity to the GlycoBind/LectinOracle training sets; the source contained 549 LectinOracle-compatible glycan tasks, and 272 exact-unseen glycans were retained. Test II retained four Sasmal AB5 proteins, with 83.1/83.1% maximum identity, from an array containing 72 unique glycans; 34 exact-unseen glycans were retained. Test III retained one EclA protein measured under two array conditions, with 3.0/3.0% maximum identity, from 487 unique glycans; 240 exact-unseen glycans were retained. Test IV retained two influenza H5 proteins and four profiles from an encoded set of 199 unique glycans, with 92.3/91.9% maximum identity; 173 exact-unseen glycans were retained. Test V retained two bisected N-glycan array proteins, with 39.5/39.1% maximum identity, from 74 array glycans; 70 exact-unseen glycans were retained. Test VI retained two proteins and six profiles from the LectinOracle main-figure subset, with 31.7/17.3% maximum identity, from 300 numeric-array glycans; 211 exact-unseen glycans were retained. None of the retained proteins had an exact sequence match to either training set.

**Figure 3.**
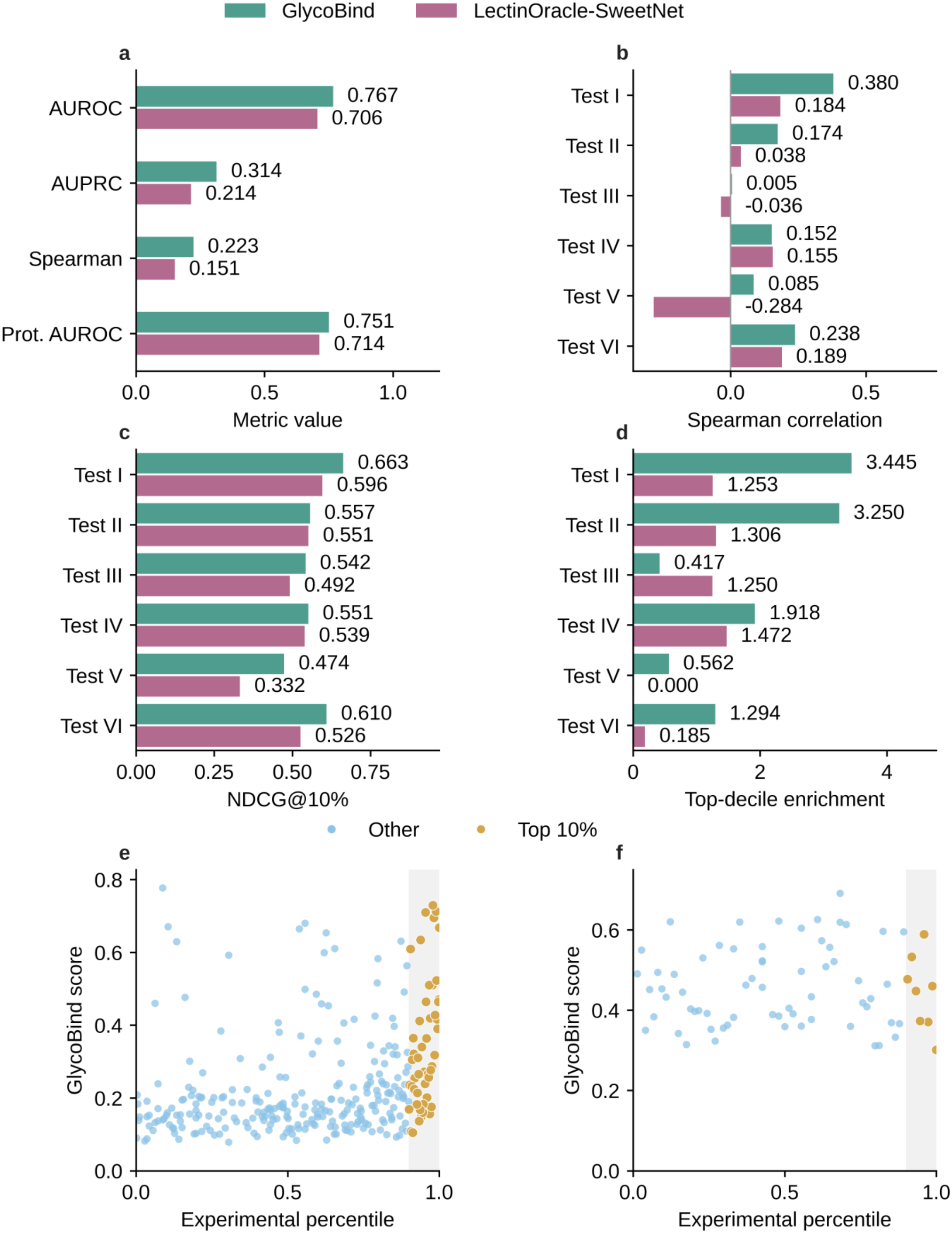
External comparison and single-profile rankings. (a) main held-out comparison with LectinOracle-SweetNet; (b) protein-profile Spearman correlation on six exact-cleaned external glycan-array datasets; (c) normalized discounted cumulative gain at the top 10% (NDCG@10%); (d) top-decile enrichment; (e) single-profile ranking for Rotavirus B VP8*B; (f) single-profile ranking for the GS-II bisected N-glycan profile. In a-d, GlycoBind and LectinOracle-SweetNet are shown as paired horizontal bars. In e and f, each point is a glycan, the horizontal axis is its experimental percentile and the vertical axis is the ensemble GlycoBind score; gold points mark the experimental top decile.

The comparator was the originally released LectinOracle-SweetNet model. For each external dataset, a task was removed when its normalized protein sequence exactly matched a GlycoBind training or validation sequence or a LectinOracle-SweetNet training sequence. A task was also removed when its canonical IUPAC glycan exactly matched a GlycoBind graph-library structure or a LectinOracle-SweetNet training structure. Comparisons used only task identifiers that both models could score. The retained benchmark contained 1,827 tasks and 19 profiles from 12 proteins. All retained glycan structures were exact-unseen.

The LectinOracle-SweetNet main-figure subset was restricted to numeric array tasks and common glycan coverage. One protein with an exact sequence match to a GlycoBind training sequence was removed. The resulting subset contained 427 tasks and six profiles.

### External ranking performance

LectinOracle-SweetNet returns a z-score for each protein-glycan pair. We therefore compared the two models by ranking rather than by raw score or probability threshold. On the main dataset, overall ranking was assessed with AUROC, AUPRC, protein-macro Spearman correlation and protein-macro AUROC. Strong-binding ranking was assessed with NDCG@10% within each protein profile, using positive labels defined by z>1.0364. GlycoBind achieved a mean NDCG@10% of 0.479±0.023, compared with 0.386±0.022 for LectinOracl e-SweetNet. Higher AUROC, AUPRC, protein-macro Spearman correlation and protein-macro AUROC were observed for GlycoBind **(Figure 3a)**.

GlycoBind achieved a dataset-balanced Spearman correlation of 0.172 versus 0.041 for LectinOracle-SweetNet. Dataset-balanced NDCG@10% was 0.566 versus 0.506, and dataset-balanced top-decile enrichment was 1.814 versus 0.911. GlycoBind achieved higher NDCG@10% in all six datasets and higher Spearman correlation and top-decile enrichment in five of six datasets; all three metrics favoured GlycoBind in four datasets. The exact-cleaned comparison is reported **(Figure 3)**. External NDCG@10% uses k = ceiling(0.10n) and continuous response percentiles as graded relevance within each profile.

The strongest external evidence came from high-intensity pair ranking. For Rotavirus B VP8*B, GlycoBind achieved Spearman 0.380 versus 0.184 and top-decile enrichment 3.445 versus 1.253. In the Sasmal AB5 panel, top-decile enrichment was 3.250 versus 1.306, and binary AUROC was 0.697 versus 0.476. Influenza H5 produced similar Spearman correlations for the two models (0.152 versus 0.155), but GlycoBind had higher NDCG@10% (0.551 versus 0.539) and higher top-decile enrichment (1.918 versus 1.472). The LectinOracle main-figure subset favoured GlycoBind, with Spearman 0.238 versus 0.189 and NDCG@10% 0.610 versus 0.526.

The difficult datasets remained visible. EclA produced near-zero Spearman correlations for both models (0.005 for GlycoBind and -0.036 for LectinOracle-SweetNet), and LectinOracle-SweetNet had higher top-decile enrichment on this dataset. The bisected N-glycan panel still favoured GlycoBind relative to LectinOracle-SweetNet, but its absolute ranking performance was lower than in the previous training-pool configuration (Spearman 0.085 and NDCG@10% 0.474). These results show that exact novelty does not guarantee useful ranking. Array dynamic range, ligand chemistry, glycan presentation, protein-family distance and the availability of informative homologues can all change external performance. Rotavirus VP8*B illustrates why a simple distance rule is insufficient; the protein and glycans were both distant, yet high-intensity pairs were strongly enriched. The external ranking results and the corresponding protein-similarity analysis are reported **(Table 2)**.

**Table 2.** External ranking performance and similarity to training data after exact cleaning.

| **Dataset** | **Proteins / glycans** | **Max identity G / L (%)** | **Spearman G** | **Spearman L** | **NDCG@10% G** | **NDCG@10% L** | **Enrichment G** | **Enrichment L** |
|---|---|---|---|---|---|---|---|---|
| Rotavirus VP8*B | 1 / 272 | 3.60 / 4.70 | 0.38 | 0.18 | 0.66 | 0.60 | 3.45 | 1.25 |
| Sasmal AB5 | 4 / 34 | 83.10 / 83.10 | 0.17 | 0.04 | 0.56 | 0.55 | 3.25 | 1.31 |
| EclA | 1 / 240 | 3.00 / 3.00 | 0.01 | -0.04 | 0.54 | 0.49 | 0.42 | 1.25 |
| Influenza H5 | 2 / 173 | 92.30 / 91.90 | 0.15 | 0.16 | 0.55 | 0.54 | 1.92 | 1.47 |
| Bisected N-glycan | 2 / 70 | 39.50 / 39.10 | 0.09 | -0.28 | 0.47 | 0.33 | 0.56 | 0.00 |
| LectinOracle main-figure | 2 / 211 | 31.70 / 17.30 | 0.24 | 0.19 | 0.61 | 0.53 | 1.29 | 0.19 |

Retained protein and glycan counts, ranking metrics and similarity to training data are reported **(Table 2)**. G denotes GlycoBind and L denotes LectinOracle-SweetNet. Protein identity is the maximum local sequence identity to the union of GlycoBind train and validation proteins across the five seeds or to LectinOracle-SweetNet training proteins. Spearman and NDCG@10% values are macro averages across profiles within each dataset; top-decile enrichment is relative to random selection. AUROC and AUPRC were available only for the binary Sasmal AB5 panel: G 0.70 and 0.65 versus L 0.48 and 0.59.

### Protein similarity audit

External protein sequences were aligned against the union of GlycoBind train and validation proteins across all five seeds and separately against the LectinOracle-SweetNet training proteins. Candidate proteins were first ranked by 5-mer containment, and the 50 nearest candidates were aligned with a local dynamic-programming aligner. Maximum identity was defined as matching aligned positions divided by the longer sequence length. Alignment coverage was normalized by the longer of the query and reference sequences.

### Single-protein glycan-array ranking

Protein-specific rankings can differ from aggregate metrics. One strongly ranked and one more weakly ranked exact-unseen example are reported **(Figure 3e,f)**. For Rotavirus B VP8*B, the GlycoBind score increased with experimental percentile across 301 exact-cleaned array entries (Spearman rho = 0.380). High experimental responses were concentrated toward the upper part of the score distribution. For the GS-II profile of the bisected N-glycan array, 72 exact-cleaned entries produced a weak Spearman correlation of 0.017 and substantial overlap between low- and high-response glycans. GlycoBind therefore ranked one distant protein-glycan profile effectively but showed limited ordering for the other exact-unseen profile.

## Conclusion and future direction

GlycoBind provides a compact and reproducible end-to-end framework for protein-glycan binding prediction from a PDB structure and an IUPAC-condensed glycan. It combines frozen DeepGlycanSite residue representations with frozen GlycanGT graph representations and a lightweight trainable head, with five-member ensembling. The model showed stable protein-held-out discrimination and enriched high-intensity binding pairs across internal and external evaluations, supporting its use for candidate prioritization before biochemical or structural follow-up.

Its applicability is strongest for protein families and glycan classes with adequate representation in the training set, particularly C-type lectin/Siglec proteins and common N- and O-glycans. Performance declines for poorly represented families, unclassified glycans and queries far from the training distribution. The model currently cannot resolve changes in protein glycan-binding profiles caused by a single-site mutation. External scores may require dataset-specific recalibration.

Future work will focus on expanding the diversity of proteins and glycans to improve model generalization, and on adopting more detailed modelling to capture how mutant proteins alter glycan recognition.

## Data and code availability

The datasets, model source code and GlycoBind toolkit are available at https://github.com/Huangxirui666/glycobind. The model release is available from the repository’s Releases page.

## Acknowledgements

This research was financially supported by National Nature Science Foundation of China (22074105) and Shanghai Science and Technology Commission (14DZ2261100).

## TOC Graphic

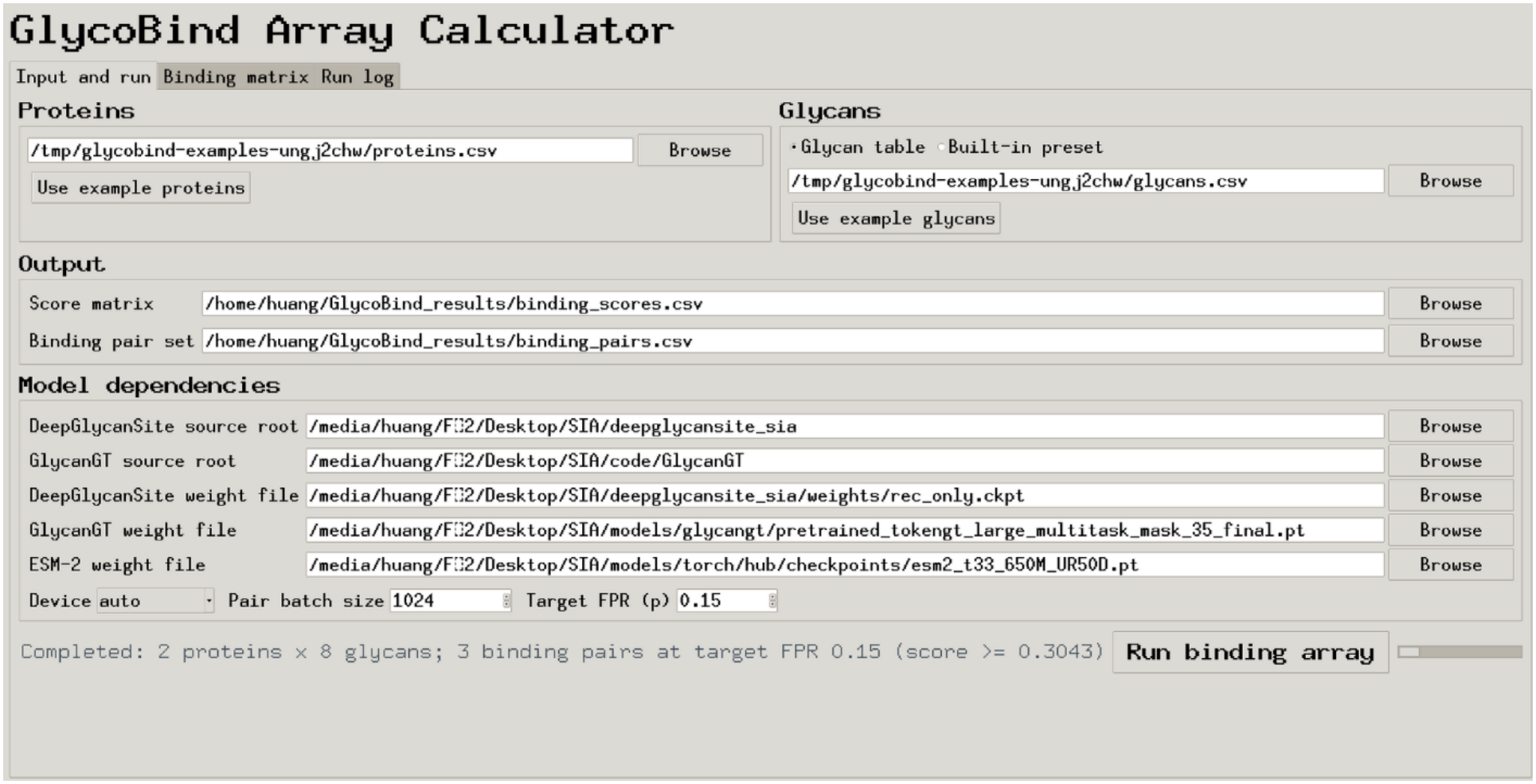

